# Structure and dynamics of the RF-amide QRFP receptor GPR103

**DOI:** 10.1101/2023.12.06.570340

**Authors:** Aika Iwama, Hiroaki Akasaka, Fumiya K. Sano, Hidetaka S. Oshima, Wataru Shihoya, Osamu Nureki

**Affiliations:** Department of Biological Sciences, Graduate School of Science, The University of Tokyo, Bunkyo, Tokyo 113-0033, Japan

## Abstract

Pyroglutamylated RF amide peptide (QRFP) is a type of peptide hormone with a C-terminal RF-amide motif. QRFP selectively activates class-A categorized GPCR, GPR103 to exert various physiological functions such as energy metabolism and appetite regulation. Here, we report the cryo-electron microscopy structure of the QRFP-GPR103-G_q_ complex at 3.3 Å resolution. Unlike class-A GPCR, QRFP adopts an extended structure baring no secondary structure, with its N-terminal and C-terminal sides recognized by extracellular and transmembrane domains, respectively, of GPR103. The C-terminal heptapeptide of QRFP penetrates into the orthosteric pocket to act in receptor activation. Particularly, the residues that recognize the RF-amide are highly conserved in the RF-amide receptors. Notably, the unique N-terminal helix-loop-helix of the receptor traps the N-terminal side of QRFP with the pendulum-like motion to guide QRFP into the ligand-binding pocket. This movement, reminiscent of class B1 GPCRs except for orientation and structure of the ligand, is critical for the high affinity binding and receptor specificity of QRFP. Structural comparisons with closely related receptors, including RY-amide peptide-recognizing GPCRs, revealed conserved and diversified peptide recognition mechanisms, providing profound insights into the biological significance of RF-amide peptides. This study not only advances our understanding of GPCR-ligand interactions, but also paves the way for the development of novel therapeutics targeting metabolic and appetite disorders and emergency medical care.

## Introduction

Neuropeptides, a diverse array of signaling molecules, orchestrate a multitude of physiological processes in organisms. Among them, neuropeptides possessing the Arg-Phe-NH_2_ (RFamide) motif at their C-termini are designated as RFamide peptides and have attracted considerable attentions because of their pivotal roles in various biological functions^1–3^. The RF-amide family encompasses five distinct peptides: Neuropeptide FF (NPFF), Prolactin-releasing peptide (PrRP), Kisspeptin (Kiss1), gonadotropin-inhibitory hormone (GnIH), and pyroglutamylated RFamide peptide (QRFP). Each peptide interacts with specific class A G-protein-coupled receptors (GPCRs), initiating a cascade of intracellular events to modulate physiological responses. Similarly, the RY-amide peptides, characterized by the signature RY motif at their C-termini, present a similar mode of action, interacting with their respective receptors to regulate physiological processes^4^. This categorization based on the C-terminal residues highlights the specificity and diversity within these peptide families. The intricate interplay between these peptides and their receptors represents a complex network critical for maintaining homeostasis and responding to environmental changes.

QRFP is a 43-amino acid RFamide peptide with a pyroglutamylated N-terminus (namely 43Rfa). QRFP was originally identified by a bioinformatics approach and reverse pharmacology^5^, and demonstrates specific activity for GPR103 with orexigenic activity^5–7^, a G_q_-coupled receptor expressed in the brain and adrenal gland. GPR103 is predominantly localized in the central nervous system, including regions such as the hypothalamus, which plays a crucial role in the regulation of energy metabolism and appetite control. QRFP and GPR103 are implicated in a variety of physiological functions ranging from the modulation of feeding behavior to the regulation of energy homeostasis^8^, cardiovascular function and bone formation^9^. This QRFP-GPR103 pair is remarkably conserved across various vertebrate species^10^, highlighting its fundamental significance in biological systems. GPR103 is not only pivotal in maintaining physiological balance but also presents therapeutic targets for disorders related to metabolism and appetite dysregulation. For example, QRFP administration reportedly increased food intake and fat mass while reducing glucose-induced insulin release, and may also cause osteopenia and facilitate nociception^11^. Thus, the GPR103 antagonist demonstrated an anorexigenic effect in mice, and has been focused as a treatment for obesity. Additionally, the GPR103 antagonist would be useful to trigger dormant state for survival treatment in emergency medical care.

Recent advancements in cryo-electron microscopy (cryo-EM) have unveiled numerous GPCR-G-protein complex structures, including those associated with C-terminally amidated peptides such as cholecystokinin, orexin, and RY-amide neuropeptide Y^12–14^. These analyses have shed light on the intricacies of ligand-receptor interactions and their activation mechanisms. Nevertheless, a notable gap remains in our understanding of RF-amide receptors, specifically GPR103. The structures of these receptors have not been elucidated, leaving the mechanisms distinguishing RF- and RY-amides enigmatic. This also includes an understanding of the selective activation of GPR103 by QRFP and the fundamental principles dictating ligand specificity among related receptors. These gaps in structural data significantly hinder the strategic development of therapeutic agents targeting GPR103. Here, we report a cryo-EM structure of the QRFP-bound GPR103•G_q_ complex, offering deep insights into its ligand recognition and selectivity, and dynamics.

## Results

### Overall structure

For the structural study, we used an N-terminally truncated form of QRFP, known as 26Rfa (Fig. 1a). 26Rfa is also found *in vivo* and possesses receptor activity, as comparable to that of 43Rfa^7,8,15^. To facilitate expression and purification, we truncated the C-terminal residues after G366 of human GPR103. We also used an engineered mini-Gα_q_ (mini-G_sqi_), a mini-G_s_ protein whose N-terminal and C-terminal residues are replaced by G_i1_ and G_q_, respectively^16^. To efficiently purify the stable GPCR-G-protein complex, the receptor and mini-G_sqi_ were incorporated in a ‘Fusion-G system’ by combining two complex stabilization techniques^17^ (Supplementary Fig. 1a, b). The modified receptor and G-protein were co-expressed in HEK293 cells and purified by Flag affinity chromatography. After an incubation with Nb35 and scFv16, which binds to mini-G_sqi_, the complex was purified by size exclusion chromatography (Supplementary Fig.1c, d). The structure of the purified complex was determined by single-particle cryo-EM analysis with an overall resolution of 3.37 Å (Table 1, Supplementary Fig. 2, and “Methods”). As the extracellular portion of the receptor was poorly resolved, we performed receptor focused refinement, yielding a density map with a nominal resolution of 3.48 Å, which was combined with the overall refined map. The resulting composite map allowed us to precisely build the atomic model of all the components, including the receptor (residues 3 to 243 and 263 to 346), ligand, G-proteins, and antibodies (Fig. 1b, c).

**Fig. 1.**
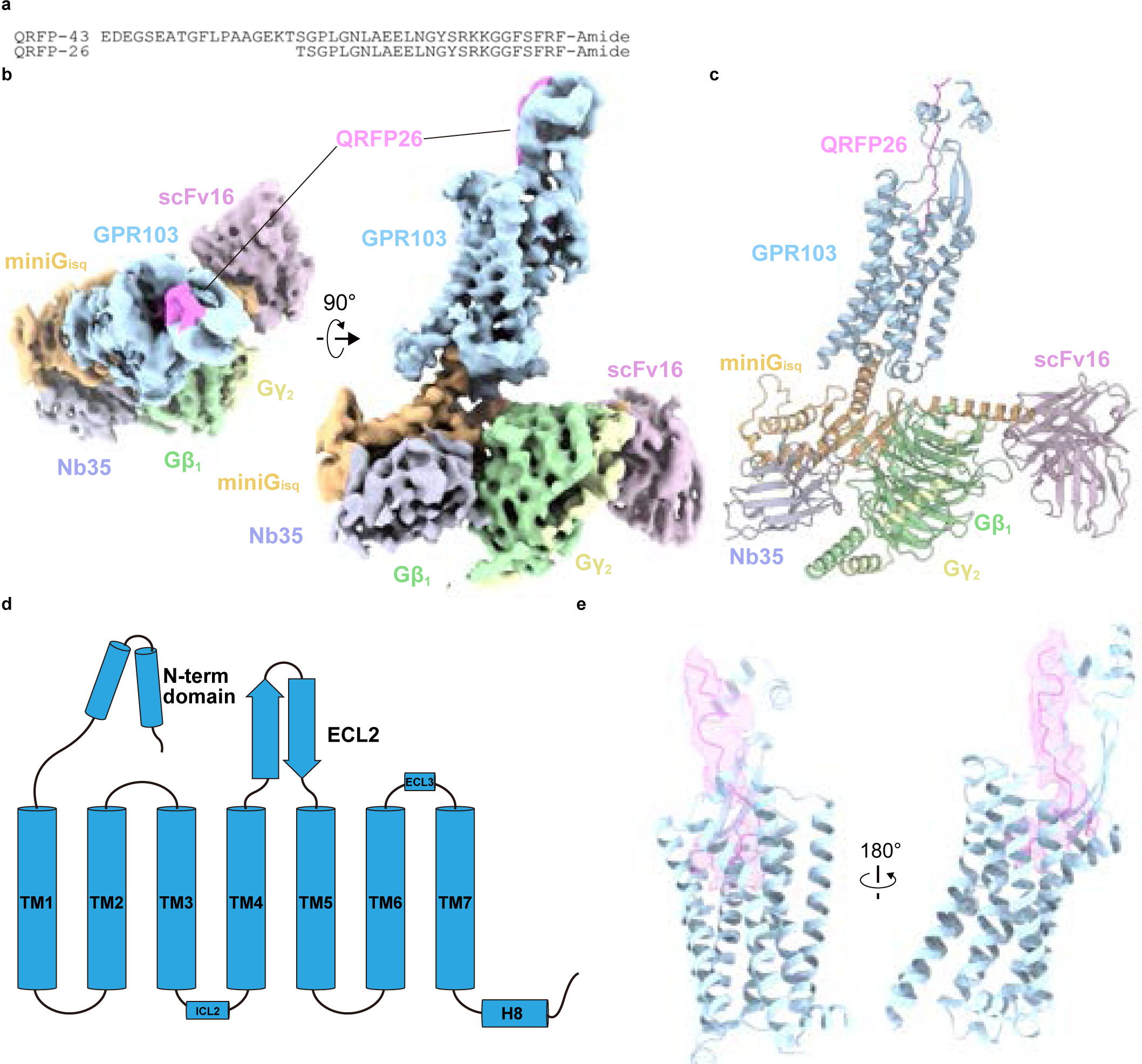
Overall structure of the GPR103-miniG_sqi_β_1_γ_2_-scFv16-Nb35 complex. **a** Amino acid sequences of QRFP43 and QRFP26. **b** Unsharpened cryo-EM density map of the GPR103-miniG_sqi_β_1_γ_2_-scFv16-Nb35 individually colored. **c** Refined structures of the complex are shown as a ribbon representation. **d** Diagram of GPR103. N-terminal forms a helix-loop-helix motif. ECL2 forms a long β sheet. **e** Ribbon representation of the QRFP26 and GPR103. Density focused on QRFP26 (pink). Two disulfide bonds are represented by stick models.

**Table. 1.** Cryo-EM data collection, refinement and validation statistics.

| Data collection | GPR103-Gq (refined) | GPR103-Gq (upright) | GPR103-Gq (tilted) |
| --- | --- | --- | --- |
| Microscope | Titan Krios (Thermo Fisher Scientific) |  |  |
| Voltage (keV) |  | 300 |  |
| Electron exposure (e<sup>-</sup>/Å<sup>2</sup>) |  | 49.6 |  |
| Detector | Gatan K3 Summit camera (Gatan) |  |  |
| Magnification |  | ×105,000 |  |
| Defocus range (μm) |  | -0.8–1.6 |  |
| Pixel size (Å/pix) |  | 0.83 |  |
| Number of movies |  | 9,555 |  |
| Symmetry |  | C1 |  |
| Picked particles |  | 11,085,317 |  |
| Final particles |  | 118,042 |  |
| Map resolution (Å) |  | 3.37 |  |
| FSC threshold |  | 0.143 |  |
| **Model refinement** |  |  |  |
| Atoms |  |  |  |
| **R.m.s. deviations from ideal** |  |  |  |
| Bond lengths (Å) | 0.003 | 0.003 | 0.003 |
| Bond angles (°) | 0.626 | 0.626 | 0.621 |
| Validation |  |  |  |
| Clashscore | 10.77 | 11.12 | 11.79 |
| Rotamers (%) | 0.1 | 0 | 0 |
| **Ramachandran plot** |  |  |  |
| Favored (%) | 96.28 | 96.36 | 95.81 |
| Allowed (%) | 3.72 | 3.64 | 4.19 |
| Outlier (%) | 0 | 0 | 0 |

The receptor consists of the canonical 7 transmembrane helices (TM) connected by three intracellular (ICL1–3) and three extracellular (ECL1–3) loops, the amphipathic helix 8 at the C-terminus (H8), and the N-terminal residues exposed on the extracellular side (Fig. 1d, e). ICL3 was disordered in the cryo-EM map. At the secondary structure level, ICL2 and ECL3 contain short α helices, and ECL2 forms a long β sheet. Two disulfide bonds are observed in the GPR103 structure (Fig. 1e). One is the highly conserved disulfide bond between C118^3.25^ (superscripts indicate Ballesteros–Weinstein numbers) and C201^ECL2^, and another unexpected disulfide bond is formed between C285^6.47^ and C327^7.48^. Notably, the N-terminal residues extend above ECL2, constituting the extracellular domain (ECD) together with ECL2. An unambiguous density was observed from the interior of the transmembrane domain (TMD) to the ECD, allowing us to model residues 7 to 26 of QRFP (Fig. 1d, e). QRFP adopts an elongated confirmation, consistent with the NMR analysis of QRFP alone^18^.

On the intracellular side, the GPR103-G_q_ complex demonstrates the typical G_q_ binding mode. Specifically, the C-terminal helix of Gα_q_ (α5h) is deeply embedded within the intracellular cavity formed by the outward displacement of TM6, resulting in the formation of an active signaling complex. In the proximity of the widely conserved N^7.49^P^7.50^xxY^7.53^ motif ^19^, the side chains of Y234^5.58^ and Y332^7.53^ are oriented towards each other (Fig. 2a). Around another conserved D^3.49^R^3.50^Y^3.51^ motif (modified to ERH in GPR103), R143^3.50^ forms a hydrogen bond with the backbone carbonyl of Y243^G.H5.23^ (superscript indicates the common Gα numbering [CGN] system^20^) in the α5h of Gα_q_. Furthermore, numerous residues in TM6 and 7 form hydrogen bonds with the C-terminus of α5h, comprising an electrostatic network (Fig. 2b). Regarding ICL2, the major interface following α5h, two remarkable interactions are observed. One is the hydrophobic interaction between ICL2 and the Gα subunit. Specifically, the bulky hydrophobic residue F151^ICL2^ is captured within a hydrophobic pocket in the Gα subunit, as in many other GPCRs^16,21^(Fig. 2c). The other is stabilization through stacking interaction: W155^ICL2^ forms π-π stacking and π-cation interactions with R32^G.hns1.03^, contributing to the stability of the active signaling complex. Finally, we compared the GPR103-G_q_ complex structure with those of other Class A GPCRs bound to various G-proteins. In terms of the binding angles for α5h, GPR103-G_q_ naturally exhibits similarity to G_q_-coupled GPCRs rather than G_s_- and G_i_-coupled GPCRs^13,16,17,22–24^(Fig. 2d, e). Overall, GPR103 demonstrates a conserved activation mechanism and G-protein binding mode, consistent with those of many class A GPCRs.

**Fig. 2.**
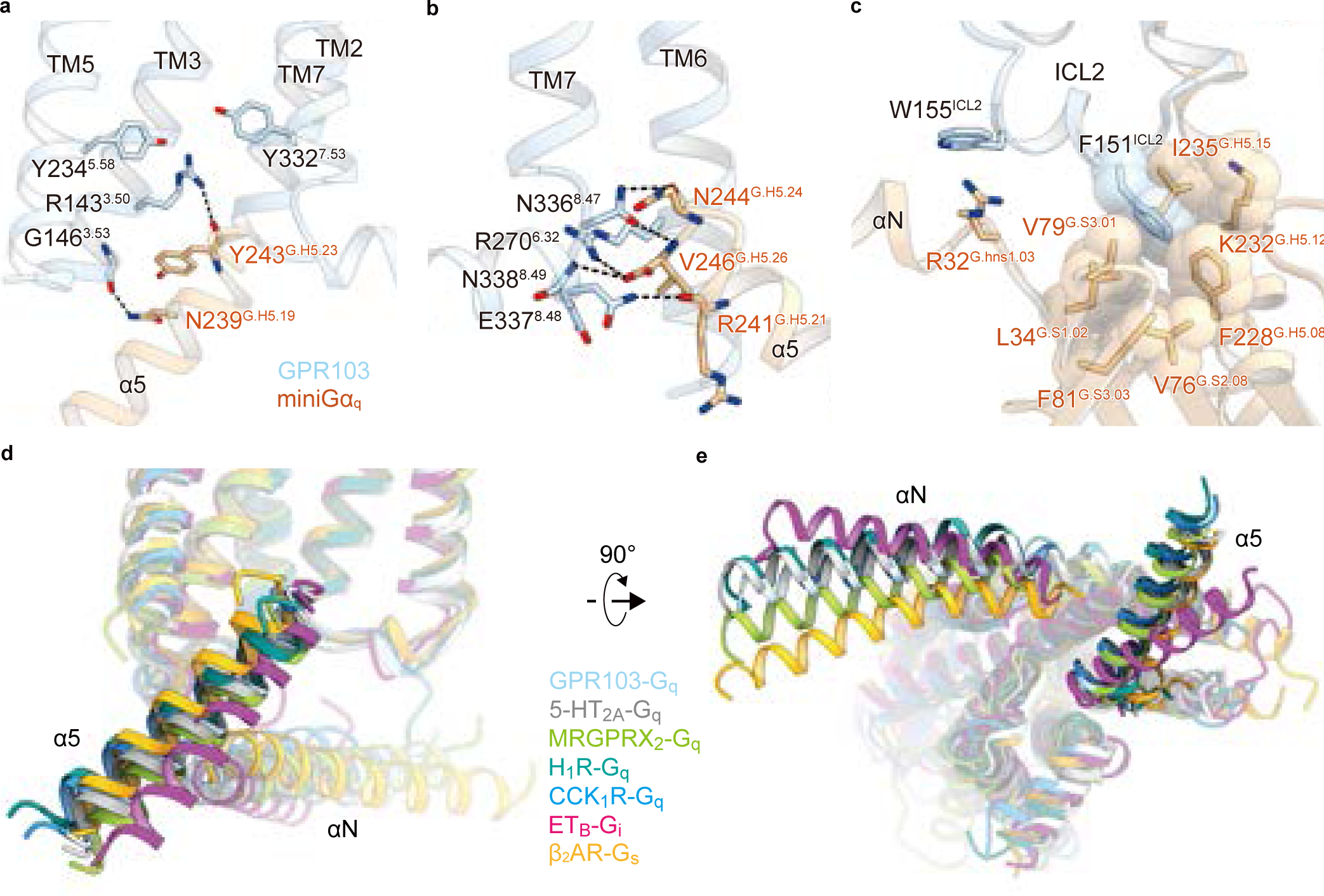
The G_q_ Binding mode. **a** Interface between GPR103 (blue) and α5h of Gα_q_ (orange) shown with critical residues and polar interactions represented by stick models and dotted lines. **b** Electrostatic network between TM6, TM7 of GPR103 and the C-terminus of α5h. **c** Interface between ICL2 of GPR103 and G_q_ shown with residues involved in hydrophobic interactions represented by CPK models. **d, e** Comparison of the angles and positions of α5h and αN relative to the receptor. G-protein bound GPCRs used in comparison are as follows: GPR103-G_q_ (blue), 5-HT_2A_-G_q_ (PDB 6WHA, gray), MRGPRX_2_-G_q_ (PDB 7S8N, yellow green), H_1_R-G_q_ (PDB 7DFL, dark cyan), CCK_1_R-G_q_ (PDB 7MBY, dodger blue), ET_B_-G_i_ (PDB 8IY5, magenta), and β_2_AR-G_s_ (PDB 3SN6, yellow orange).

### QRFP binding site in the transmembrane domain

QRFP binds to both the ECD and TMD with its C-terminal amide directed toward the TMD core (Fig. 3a) The C-terminal heptapeptide of QRFP, fits vertically within the TMD and creates an extensive interaction network with TMs 2-7 and ECL2 of the receptor (Fig. 3b, c, Supplementary Fig. 3). This interaction between QRFP and the TMD can be broadly divided into those with RF-amide and the rest (residues 20–24). In the latter, F22 and F24 of QRFF form extensive hydrophobic interactions with the extracellular halves of TM2 and TM3, leading to the upright structure of QRFP inside the TMD (Fig. 3c). In addition, several hydrogen bonds exist between the receptor and the peptide backbone of QRFP. In comparison, RF-amide binds very tightly deep in the pocket (Fig. 3b), consistent with its significance in receptor binding reported in previous studies with mutant peptides. Specifically, the C-terminal amide forms hydrogen bonding interactions with T102^2.61^, Q125^3.32^, and Q318^7.39^. The side chain of F26 is present at the deepest position in the TMD, and surrounded by bulky hydrophobic residues in TM2, TM3, and TM6. The R25 residue forms electrostatic interactions with E203^ECL2^ and E297^6.59^, in addition to a hydrogen-bond with T215^5.39^. These two residues make the ligand-binding pocket of the TMD negatively charged (Fig. 3a). Thus, the C-terminal amidation functions not only in interactions with the receptor but also in enhancing the charge complementarity with the pocket, by neutralizing the negative charge of the C-terminal carboxylate.

**Fig. 3.**
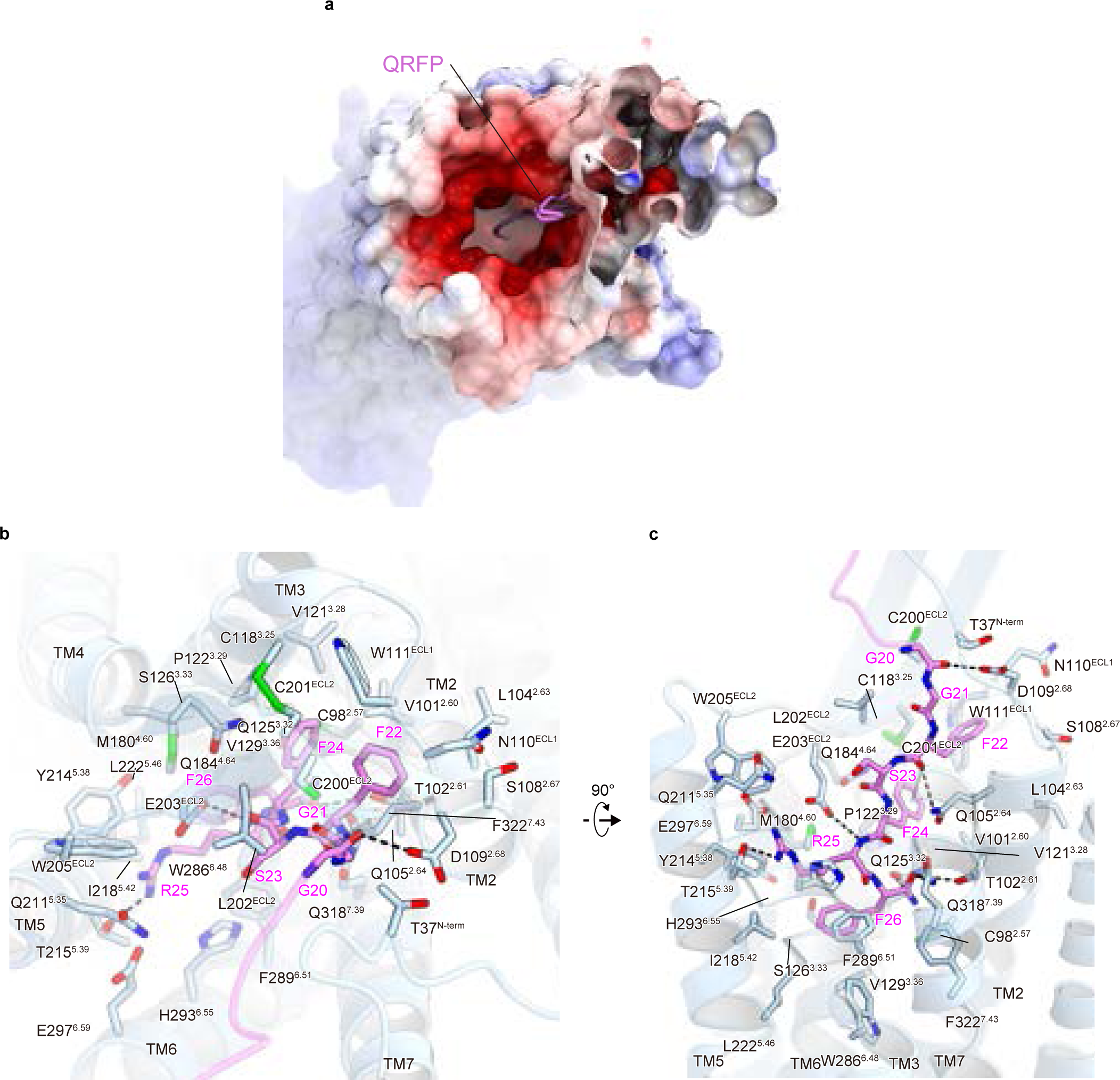
QRFP binding site in the transmembrane region. **a** Surface representation of GPR103 viewed from the extracellular side. Positive and negative charges of the receptor are colored in blue and red, respectively. **b, c** Binding pocket for QRFP in the TMD, viewed from the extracellular side **(b)** and membrane plane **(c)**. Residues involved in the QRFP-GPR103 interaction within 4.5 Å are shown as pink and blue sticks, respectively. Black dashed lines indicate hydrogen bonds.

Previous studies with mutant peptides have shown that the C-terminal heptapeptide is sufficient to activate GPR103^25,26^, despite its reduced affinity. The heptapeptide is conserved from fish to mammals, except for S23(Supplementary Fig. 4a), consistent with the fact that the S23 side chain poorly interacts with the receptor. The receptor residues interacting with the heptapeptide are highly conserved among the homologs (Supplementary Fig. 4b, c). Accordingly, this observed heptapeptide-TMD interaction plays a key role in evolutionarily-conserved GPR103 activation.

This RF amide is the only conserved part of the RF amide peptide (Supplementary Fig. 5a). To examine the conservation of its recognition mechanism, we compared the residues interacting with the RF amide with the corresponding RF amide receptors (GPR10, GPR54, GPR74, and GPR147) (Supplementary Fig. 5b, c). T102^2.61^ and Q125^3.32^, which recognize the C-terminal amide of QRFP, are completely conserved. Although Q318^7.39^ is replaced by histidine in all the other RF-amide receptors, it would form hydrogen bonds with the oxygen atom of the C-terminal amide of QRFP. These considerations suggest that amide recognition by hydrogen bonding interactions via these three residues is a common mechanism in RF-amide receptors. The two negative charges near R25 are also conserved in RF-amide receptors, although E297^6.59^ is replaced by alanine in GPR54. Instead, Q211^5.35^, which forms a polar contact with R25 in GPR103, is replaced by E201^5.35^ in GPR54, suggesting the conserved recognition of R25 by the two negative charges. In contrast, the recognition of the F26 side chain is less stringent but is shared by a bulky hydrophobic amino acid. Thus, the RF amide recognition mechanism observed in QRFP-GPR103 is highly conserved in RF amide receptors.

### Unique architecture of the N-terminal region

We observed an unambiguous density above ECL2, despite the suboptimal local resolution ranging from 4 to 6 Å (Fig. 4a and Supplementary Fig. 2). The density aligned with the N-terminal structure was predicted by AlphaFold^27–29^. To enhance the fidelity of the map, we performed a 3D flexible refinement implemented in cryoSPARC^30^(Fig. 4b and Supplementary Fig. 6a-f). This improved map facilitated the accurate model building of the predicted N-terminal structure onto the density, employing both rigid body and all-atom refinement implemented in COOT. Furthermore, we successfully modeled QRFP26 up to N7. Thus, the current model allowed us to discuss the secondary structures of the N-terminal residues.

**Fig. 4.**
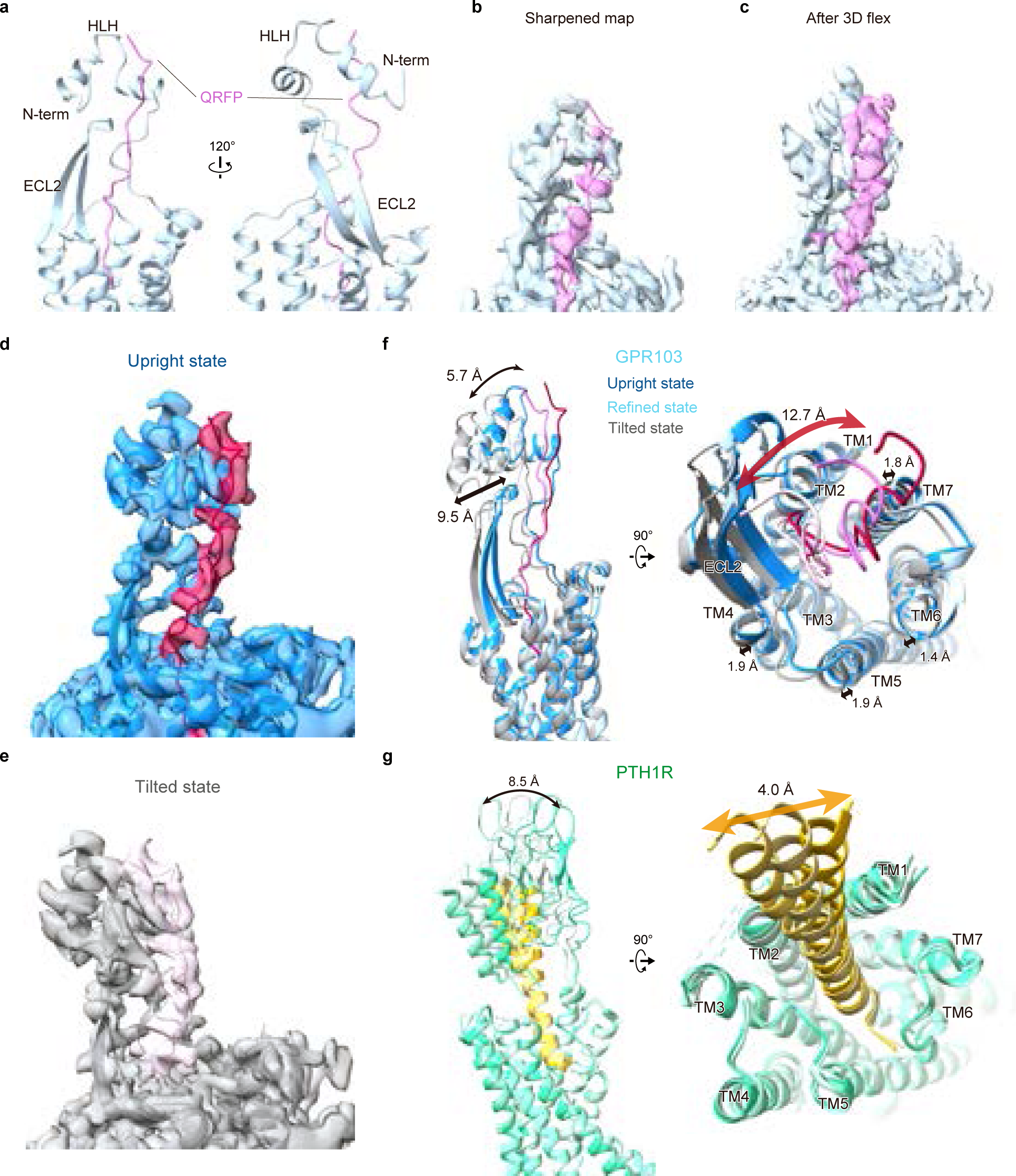
Unique architecture of the extracellular region. **a** Ribbon representation of GPR103 focused on the extracellular side. **b** Model and density map sharpened by phenix.auto_sharpen. **c** Model and density map after 3D flexible refinement. **d, e** The two most significantly changed maps by 3D flexible refinement. **f, g** Superpositions of ECD changes in GPR103 **(f)**, and PTH1R (PDB 7vvm, 7vvl, and 7vvk) **(g)**.

The N-terminal residues, numbering 2 through 40, of GPR103 extend from TM1 to above ECL2 and form a helix-loop-helix (HLH) (Fig. 4c). This N-terminal HLH motif is conserved among GPR103 homologs (Supplementary Fig. 4c), and our homology searches have failed to identify any similar sequences in other proteins, indicating unique feature of GPR103. QRFP extends vertically from the G20, interacting predominantly with the ECD and becoming sandwiched by the N-terminal HLH of the receptor. While QRFP20–26 are sufficient for receptor activation, their EC_50_ value is 75-fold lower compared to QRFP26^25,26^. This fact suggests that the ECD functions as an affinity trap, achieving high compatibility through interactions with the N-terminal residues of QRFP26.

The 3D flexible refinement also uncovered a significant conformational alteration on the extracellular side (Supplementary Fig. 6g, h and Supplementary Movie 1). We then built the models of this alteration on the two most significantly changed maps among the output (Fig. 4d, e and Table 1). The results revealed the upright and tilted states of the ECD. A structural comparison of the two states elucidated the dynamic movement of the N-terminal HLH by about 10 Å (Fig. 4f). Moreover, QRFP26, the ECD, and the extracellular half of the TMD oscillate like a pendulum with the C-terminus of QRFP26 as the base point. In the original map, the ECD of the refined structure is positioned in between the upright and tilted states, whereas the transmembrane helices are more closely aligned with the upright, implying the fundamental stability of the upright state.

To our knowledge, this N-terminal ECD configuration has never been observed in the class A GPCR structures or the AlphaFold database^29^, including the other RF-amide receptors. Moreover, among the RF amides other than QRFP, three consist of fewer than 12 residues, and the longer PrRP-31 lacks sequence homology except for the C-terminal RF-amide (Supplementary Fig. 5a). Consequently, the structure and dynamics of the ECD and QRFP observed in this study are unique to GPR103, thus playing a critical role in the GPR103 selectivity of QRFP.

Other classes of GPCRs commonly possess N-terminal domains with various lengths and functional roles, depending on the class. For example, class B GPCRs, akin to GPR103 as peptide receptors, feature an ECD with a ∼150 amino acid ’hotdog-like’ domain that acts as an affinity trap by encasing the agonist peptide^31–33^(Fig. 4g). Despite significant differences in sequence homology and length, a functional analogy exists between the ECDs of GPR103 and class B1 GPCRs. Additionally, parathyroid hormone receptor 1 (PTH1R) in class B1 GPCRs exhibits structural polymorphism in its ECD^32^ (Fig. 4g), similar to the fluctuating actions observed in GPR103 (Fig. 4f). However, only the N-terminal ECD of PTH1R moves independently of the TMD, in stark contrast to GPR103 where the entire ligand-binding pocket, iincluding the ECD, undergoes structural changes. This distinction may be attributed to the ligand conformation: class B1 GPCR ligands form a rigid, straight helical structure, whereas QRFP does not adopt any secondary structure.

### Structural comparison with related peptide receptors

GPR103 exhibits a high degree of sequence homology with CCK receptors (CCKRs), orexin receptors (OXRs), and RY-amide neuropeptide Y receptors (YRs), which uniformly recognize peptides with amidated C-termini (Supplementary Fig. 5a) via their TMDs. To elucidate the characteristics of the structure and peptide recognition mechanisms of GPR103, we compared the structures of Y_1_R, CCK_1_R, and OX_2_R in complex with G_q_ ^12–14^ (Fig. 5a). The lengths and secondary structures of the peptide ligands are diverse, and correspondingly, the lengths of the β-sheet in ECL2 are different. This comparison further highlights the distinctiveness of the N-terminal structure of GPR103 for peptide recognition. Despite the differences on the extracellular side, the C-termini of the agonist peptides as well as the intracellular sides of the receptors, superimposed well, suggesting an evolutionary linkage among these receptors.

**Fig. 5.**
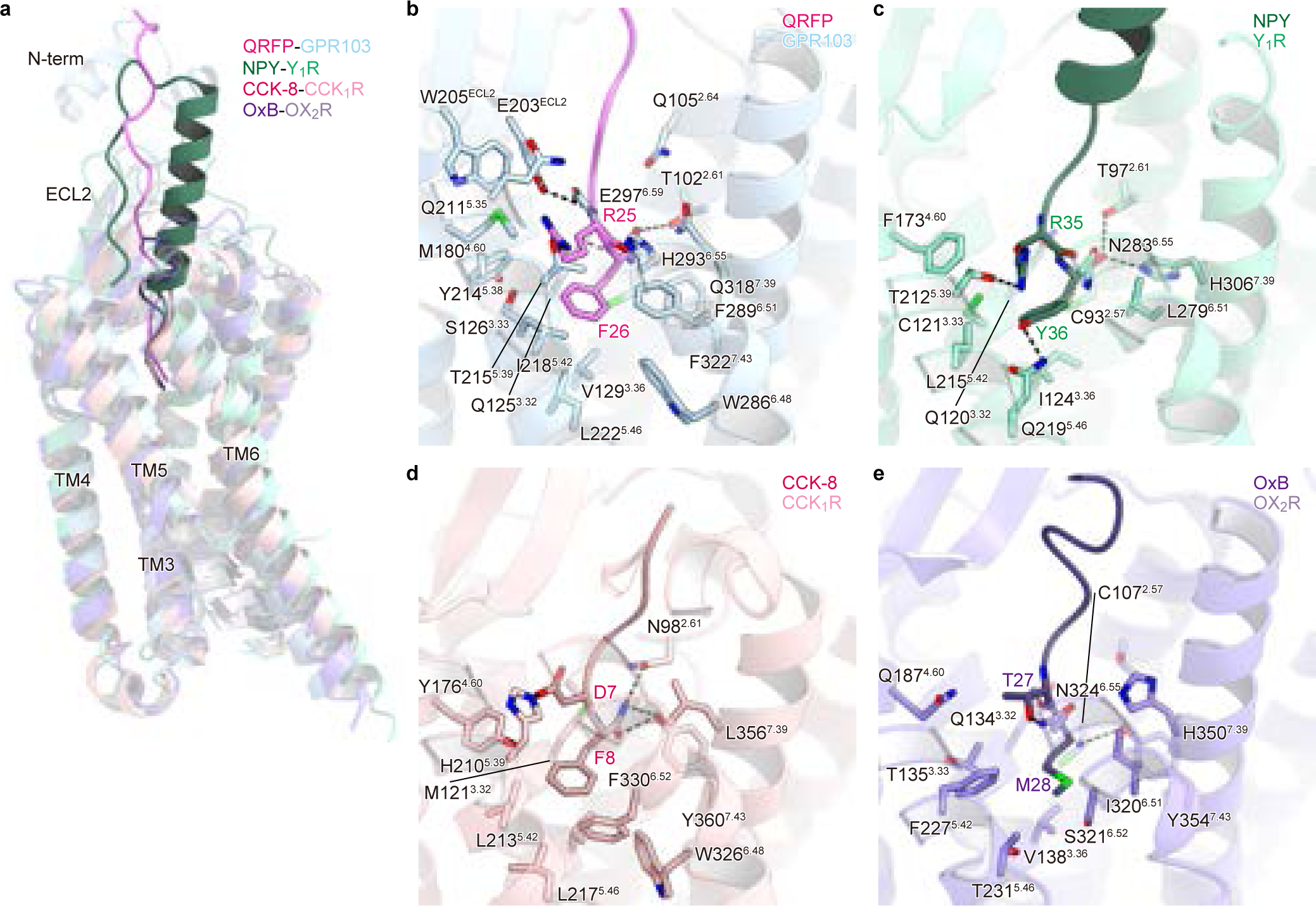
Comparison of other related amide-peptide receptors. **a** Superimposition of GPR103 (blue), Y_1_R (PDB 7X9A, green), CCK_1_R (PDB 7MBY, magenta), and OX_2_R (PDB 7L1U, purple) structures in complex with QRFP26, NPY CCK-8, and OxB, peptides, respectively. **b-e** Binding mode of the peptide ligands within the transmembrane domain of GPR103 **(b)**, Y_1_R **(c)**, CCK_1_R **(d)**, and OX_2_R **(e)**. The residues involved in ligand-receptor interactions are represented by stick models. Black dashed lines indicate hydrogen bonds.

We also examined the interactions of the C-terminal residues of the ligand in GPR103 and Y_1_R (Fig. 5b, c), representing the RF and RY amide receptors, respectively. In both instances, the C-terminal arginine of the ligand forms a hydrogen bond with T^5.39^ and ionic interactions with either aspartate or glutamate. The carbonyl oxygen of the C-terminal amide hydrogen bonds with T^2.61^ and Q/H^7.39^. The nitrogen of the amide hydrogen bonds with Q^3.32^, with C^2.57^ in close vicinity. In Y_1_R, the nitrogen and C93^2.57^ are uniquely positioned to form a hydrogen bond (Fig. 5c), a feature not found in GPR103 (Fig. 5b), although this does not preclude the potential for such interactions *in vivo*. These highly conserved residues within the RF and RY amide receptors imply a shared recognition mechanism for the C-termini of the peptide ligands.

Next, we focused on the difference between the recognition of the RY-amide and RF-amide. In Y_1_R, Q219^5.46^ forms a hydrogen bond with the hydroxyl group of the C-terminal Y36 of the peptide ligand (Fig. 5c), which is highly conserved in the NPY receptors (Supplementary Fig. 5b). Exceptionally, Q219^5.46^ is replaced by L^5.46^ in Y_2_R^34^(Supplementary Fig. 5b), but its cryo-EM structure revealed that S^5.42^, situated one turn above, alternatively forms a hydrogen bond with Y36. Contrarily, in the QRFP-GPR103 complex, the hydrogen bond between Y36 and Q219^5.46^ is replaced by a hydrophobic interaction between F26 and L222^5.46^. L^5.46^ is highly conserved in the RF-amide receptors (Supplementary Fig. 5b). Exceptionally, in GPR10 L^5.46^ is replaced by the polar residue T^5.46^, which is not expected to interact with the C-terminal phenylalanine, due to its shorter side chain. The presence or absence of a residue capable of hydrogen bonding with the hydroxyl group of tyrosine at the C-terminus may distinguish the RF and RY amide receptors.

We also compared the interactions of the C-terminal two residues of GPR103 with CCK_1_R and OX_2_R (Fig. 5d, e). In CCK_1_R, the amidated end of the peptide forms a hydrogen bond with N98^2.61^, similar to T102^2.61^ in GPR103. Furthermore, in CCK_1_R and OX_2_R, the amide group hydrogen bonds with Y^7.43^. Thus, while the amide recognition mechanism is conserved in CCK_1_R and GPR103 and in CCK_1_R and OX_2_R, GPR103 and OX_2_R are markedly different. In CCK8, the C-terminal F8 of the ligand is encased in a hydrophobic pocket similar to QRFP, whereas D7 forms a salt bridge with H210^5.39^, instead of the hydrogen bond between T215^5.39^ and R25 of QRFP (Fig. 5b, d). Meanwhile, the C-terminal M28 of OxB only contacts a few hydrophobic residues, and T27 forms a hydrogen bond with N324^6.55^, instead of T215^5.39^ in GPR103 (Fig. 5b, e). Thus, in addition to the marked differences in the extracellular region, two important residues characterized by the ligands are recognized by different mechanisms in the TMD and are crucial for the selective acceptance of the respective ligand peptides.

## Discussion

Our study has revealed the binding mode of the C-terminal QRFP heptapeptide to the TMD, which is sufficient for the activation of GPR103. The conserved recognition mechanism of the C-terminal RF-amide in various RF-amide receptors suggests broader biological significance. Comparisons with other evolutionarily close peptide receptors demonstrated the commonality and diversity in the recognition mechanisms of the C-terminal two residues of peptide ligands. In particular, the presence or absence of residues that can form hydrogen bonds with the C-terminal side chain determines whether they function as RF- or RY-amide receptors. We identified the N-terminal HLH structure of GPR103, which captures the N-terminal side of QRFP and is quite unique compared to class A GPCRs. We observed a pendulum-like motion of the ECD, including the N-terminus and the entire pocket, reminiscent of class B1 GPCRs. It should be noted that orientation and secondary structure of the peptide ligands totally differ, since the C-terminus of the α-helical ligand binds the extracellular domain in class B1 GPCRs. The structure and dynamics of these extracellular regions are important for promoting the specific, high-affinity binding of QRFP to GPR103. The distinctive structure and function of GPR103 determined in this study will be useful in the design of potential therapeutics targeting GPR103 for energy metabolism and appetite regulation.

Although the current active conformation was modeled based on the AlphaFold2 predicted structure^29^, there are considerable differences between them. Notably, the intracellular side of TM6 is in a closed configuration (Supplementary Fig. 7a, b), and R143^3.50^ of the D^3.49^R^3.50^Y^3.51^ motif (modified to E^3.49^R^3.50^H^3.51^ in GPR103) forms a salt bridge with E142^3.49^ and is not oriented toward the intracellular face (Supplementary Fig. 7c), indicating that the predicted conformation represents the inactive state. This observation allowed us to discuss the mechanism of receptor activation upon QRFP binding, by comparing the cryo-EM structure with the predicted structure. Within the ligand-binding pocket in the TMD, significant conformational changes were observed in TM5 and TM6 (Supplementary Fig. 7d, e). The hydrogen-bonding interaction between R25 and T215^5.39^ displaces TM5 inward by 1.4 Å. Conversely, F26 in QRFP induces the steric outward movement of F289^6.51^, resulting in the outward displacement of TM6 by 1.6 Å. This movement of F26 drives W286^6.48^ downward, thereby exerting a downward force on F282^6.44^ in the underlying P^5.50^-I^3.40^-F^6.44^ motif. This action is an important component of the P^5.50^-I^3.40^-F^6.44^ motif reorganization frequently observed in class A GPCRs^19^ and leads to the opening of the intracellular machinery of TM6. Such receptor activation, characterized by conformational changes in W286^6.48^, is a widespread phenomenon in class A GPCRs (e.g., human endothelin ET_B_ receptor)^16,17,33,35–44^.

In summary, these insights offer a comprehensive understanding of the mechanism of GPR103 activation upon QRFP binding. In the apo state, the N-terminal structure is hypothesized to be more labile than observed (Fig. 6a), due to its instability with merely two helices and a lower AlphaFold predictive score^29^. QRFP may initially interact with either the ECD or TMD (Fig. 6b). Eventually, QRFP establishes stable binding to both domains, presumably maintaining the interaction of the C-terminus with the receptor while allowing for some fluctuation in the ECD and ligand binding pocket in the TMD (Fig. 6c). This binding would reorganize the aromatic residue cluster in TM6, leading to the intracellular opening for receptor activation.

**Fig. 6.**
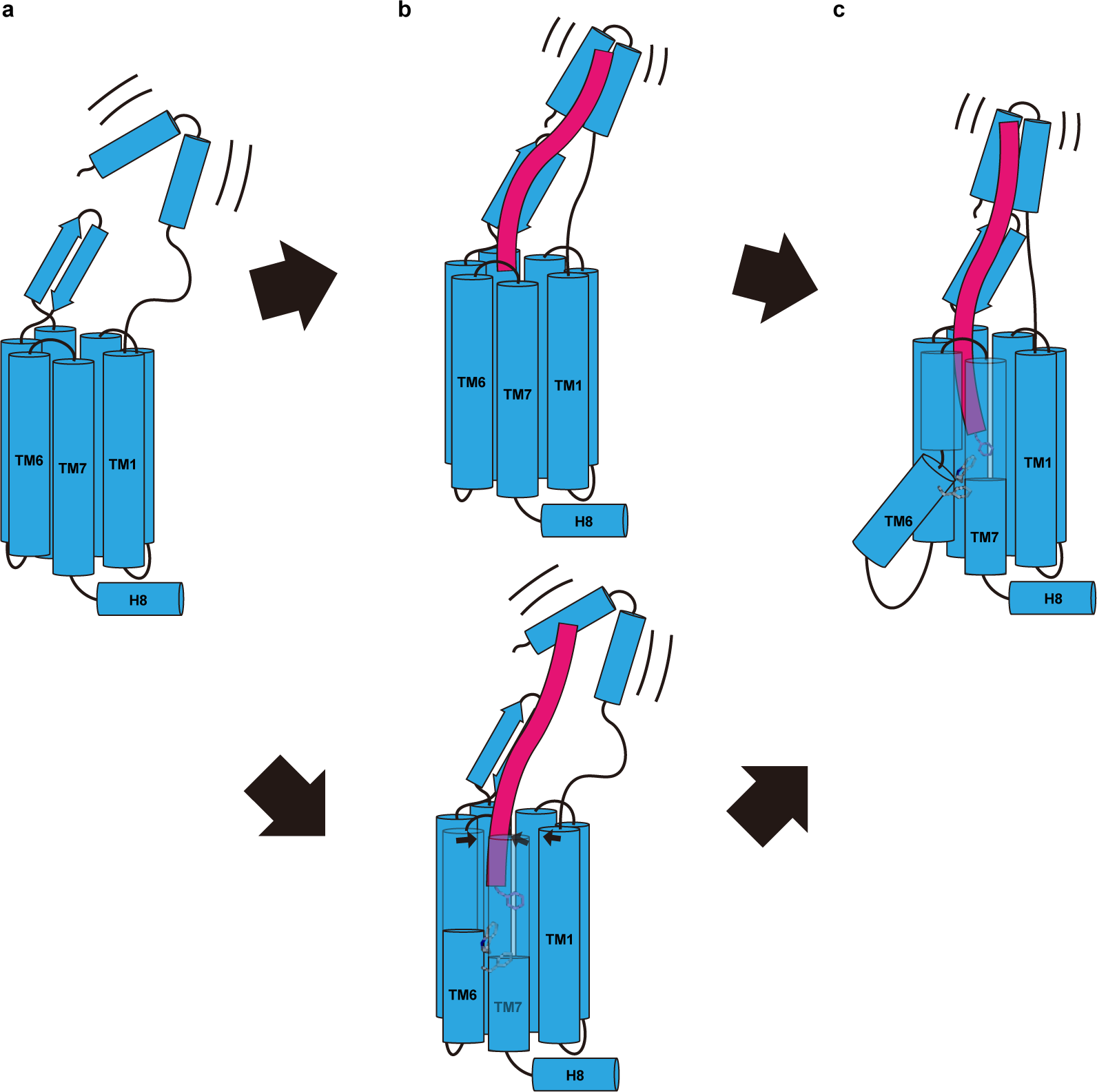
Working model for receptor activation. **a-c** Schematic representations of GPR103 activation model upon QRFP binding. From the apo state **(a)**, the peptide binds **(b)** and changes to the active state **(c)**.

In the current study, we have not only elucidated the architecture of the receptor but also shed light on the structure of QRFP itself, potentially paving the way for the modulation of QRFP-neurons (Q-neurons)^45^. QRFP is a biomarker for neurons and is capable of inducing an extended state of hypothermia and reduced metabolism, akin to hibernation. The QRFP-induced hibernation-like state (QIH) provides a unique model to study metabolic suppression for therapeutic purposes, as well as artificial hibernation. QIH through pharmacogenetics^46^ or optogenetics^47^ is achievable in murine models, but the prospect of non-genetically triggering these neurons might represent a groundbreaking stride in the realm of synthetic hibernation research. Nevertheless, Q-neurons constitute a remarkably limited subset, and currently there are no established methodologies for their selective pharmacological activation. Among the myriad strategies proposed, one promising avenue is the engineering of synthetic low-molecular-weight binders targeting QRFP, with their integration into novel drug delivery systems. The field of protein design is undergoing a rapid metamorphosis, fueled by recent innovations in structural prediction via AlphaFold^29^ and the utilization of generative AI^45^. This is exemplified by the development of soluble GPCR variants and binders based on GPCR structures. Harnessing our structural insights to conceive binders that specifically target QRFP could facilitate studies of Q-neurons. Although this idea is still in its nascent stage, it harbors substantial promise for future applications, e.g. in emergency medical care.

### Supplementary Movie 1 3D flex motion

A representation of flexibility for QRFP and ECD of QRFPR generated by 3DFlex.

## Methods

### Expression and purification of scFv16 and Nb35

The gene encoding scFv16 was synthesized (GeneArt) and subcloned into a modified pFastBac vector^48^, with the resulting construct encoding the GP67 secretion signal sequence at the N terminus, and a His_8_ tag followed by a TEV cleavage site at the C terminus. The His_8_-tagged scFv16 was expressed and secreted by Sf9 insect cells, as previously reported^48^. The Sf9 cells were removed by centrifugation at 5,000g for 10 min, and the secreta-containing supernatant was combined with 5 mM CaCl_2_, 1 mM NiCl_2_, 20 mM HEPES (pH 8.0), and 150 mM NaCl. The supernatant was mixed with Ni Superflow resin (GE Healthcare Life Sciences) and stirred for 1 h at 4 °C. The collected resin was washed with buffer containing 20 mM HEPES (pH 8.0), 500 mM NaCl and 20 mM imidazole, and further washed with 10 column volumes of buffer containing 20 mM HEPES (pH 8.0), 500 mM NaCl and 20 mM imidazole. Next, the protein was eluted with 20 mM Tris (pH 8.0), 500 mM NaCl and 400 mM imidazole. The eluted fraction was concentrated and loaded onto a Superdex200 10/300 Increase size-exclusion column, equilibrated in buffer containing 20 mM Tris (pH 8.0) and 150 mM NaCl. Peak fractions were pooled, concentrated to 5 mg/ml using a centrifugal filter device (Millipore 10 kDa MW cutoff), and frozen in liquid nitrogen.

Nb35 was prepared as previously reported^49,50^. In brief, Nb35 was expressed in the periplasm of *E. coli*. The harvested cells were disrupted by sonication. Nb35 was purified by nickel affinity chromatography, followed by gel-filtration chromatography, and frozen in liquid nitrogen.

### Constructs for expression of GPR103 and G_q_

The human GPR103 gene (UniProtKB, Q96P65) was subcloned into a modified pFastBac vector^17^, with an N-terminal haemagglutinin signal peptide followed by the Flag-tag epitope (DYKDDDDK) and the LgBiT fused to its C-terminus followed by a 3 C protease site and EGFP-His_8_ tag. A 15 amino sequence of GGSGGGGSGGSSSGG was inserted into both the N-terminal and C-terminal sides of LgBiT. Rat Gβ_1_ and bovine Gγ_2_ were subcloned into the pFastBac Dual vector. In detail, rat Gβ_1_ was cloned with a C-terminal HiBiT connected with a 15 amino sequence of GGSGGGGSGGSSSGG. Moreover, mini-G_sqi_ was subcloned into the C-terminus of the bovine Gγ_2_ with a nine amino sequence GSAGSAGSA linker. The resulting pFastBac dual vector can express the mini-G_sqi_ trimer.

### Expression and purification of the human GPR103 – G_q_ complex

The recombinant baculovirus was prepared using the Bac-to-Bac baculovirus expression system (Thermo Fisher Scientific). For expression, 0.8 L of HEK293S cells at a density of 3 × 10^6^ cells/mL were co-infected with baculovirus encoding GPR103 and miniGsqi trimer at the ratio of 2:1. Twenty hours after infection, 10 mM of Sodium Butyrate was added, and the cells were incubated at 30℃. After 48 hours, the collected cells were resuspended and dounce-homogenized in 20 mM Tris-HCl, pH 8.0(4℃), 150 mM NaCl, 10% Glycerol, 4 μM QRFP26, 5.2 μg/mL aprotinin, 2.0 μg/ml leupeptin, and 100 μM PMSF. After homogenization, Apyrase was added to the lysis at a final concentration of 25 mU/ml and the lysate was incubated at room temperature for 1 h. The crude membrane fraction was collected by ultracentrifugation at 180,000*g* for 1 h and solubilized in buffer, containing 50 mM Tris-HCl, pH 8.0, 150 mM NaCl, 1.5% Lauryl Maltose Neopentyl Glycol (LMNG) (Anatrace), 0.15 % 0.2% cholesteryl hemisuccinate (CHS) (Merck), 10% glycerol, 5.2 μg/ml aprotinin, 2.0 μg/ml leupeptin, and 100 μM PMSF, 25 mU/ml Apyrase, 4 μM QRFP26 for 2 h at 4 °C. The supernatant was separated from the insoluble material by ultracentrifugation at 180,000*g* for 30 min and incubated with 4 mL of Anti-DYKDDDDK G1 resin (Genscript) for 1 hour at 4℃. The resin was washed with 20 column volumes of buffer containing 20 mM Tris-HCl, pH 8.0 500 mM NaCl, 10% Glycerol, 0.01% LMNG, 0.001% CHS and 0.1 μM QRFP26. The complex was eluted in buffer containing 20 mM Tris-HCl, pH 8.0, 150 mM NaCl, 0.01% LMNG, 0.001% CHS, 10 μM QRFP26 and 0.2 mg/mL Flag peptide. The eluate was incubated with the Nb35 and scFv16 at 4 ℃. The complex was concentrated and purified by size exclusion chromatography on a Superose 6 increase (GE) column in 20 mM Tris-HCl, pH8.0, 150 mM NaCl, 0.01% LMNG, 0.001% CHS and 0.1 μM QRFP26. QRFP26 was added to the peak fraction to the final concentration of 4 µM and concentrated to 13.8 mg/ml.

### Sample vitrification and cryo-EM data acquisition

The purified complex was applied onto a freshly glow-discharged Quantifoil UltraAu grid (R1.2/1.3, 300 mesh), and plunge-frozen in liquid ethane by using a Vitrobot Mark IV. Data collections were performed on a 300kV Titan Krios G3i microscope (Thermo Fisher Scientific) and equipped with a BioQuantum K3 imaging filter and a K3 direct electron detector (Gatan).

First, 9555 movies were acquired with a calibrated pixel size of 0.83 Å pix^-1^ and with a defocus range of -0.8 to -1.6 μm, using EPU. Each movie was acquired for 2.6 s and split into 48 frames, resulting in an accumulated exposure of about 49.6 electrons per Å^2^ at the grid.

### Image processing

All acquired movies were dose-fractionated and subjected to beam-induced motion correction implemented in RELION 3.1^51^. The contrast transfer function (CTF) parameters were estimated using CTFFIND 4.0^52^. A total of 11,085,317 particles were extracted. The particles were subjected to 2D classifications, Ab-initio reconstruction and several rounds of hetero refinement in cryoSPARC^53^. Next, the particles were re-extracted and exported to RELION 3.1^54^, then subjected to 3D classification with a mask on the receptor. Then, the particles were subjected to Bayesian polishing in RELION. The particle sets were exported to cryoSPARC, subjected to CTF refinement, and Non-uniform refinement, yielding a map with a global nominal resolution of 3.37 Å, with the gold standard Fourier Shell Correlation (FSC=0.143) criteria^55^. Moreover, the 3D model was refined with a mask on the receptor. As a result, the receptor has a higher resolution with a nominal resolution of 3.48 Å. The overall and receptor focused maps were combined by phenix^56^. The processing strategy is described in Supplementary Figure 2.

To investigate flexibility of QRFP and ECD of QRFPR, the batch of particles was further processed by 3D flexible refinement^30^. After an overall non-uniform refinement, the mesh was prepared using micelle removed density map. Then, following the 3D flex train job and the reconstruction job, the flexibility of the QRFP and the ECD of QRFPR were visualized by the flex generate job. The processing strategy is described in Supplementary Figure 6.

### Model building and refinement

The density map was sharpened by phenix.auto_sharpen^57^ and the quality of the density map was sufficient to build a model manually in COOT^58,59^. The model building was facilitated by the Alphafold-predicted structure and cryo-EM structure of OX_2_R (PDB 7LIU)^12^. We manually fitted GPR103, the G_q_ heterotrimer and scFv16 into the map. We then manually readjusted the model into using COOT and refined it using phenix.real_space_refine^56,60^ (v.1.19) with the secondary-structure restraints using phenix secondary_structure_restraints. For the densities derived from 3DFlex, the rigid body refinement and all atom refinement were performed on two representative frames from the default 40 frames using the model without 3D flex.

### Data Availability

The cryo-EM density map and atomic coordinates for the LPA_1_-G_i_ complex have been deposited in the Electron Microscopy Data Bank and the PDB, under accession codes: EMD-XXXXX [https://www.ebi.ac.uk/emdb/entry/EMD-XXXXX], and PDB XXXX [http://doi.org/10.2210/pdbXXXX/pdb]. Source data are provided with this paper. All other data are available from the corresponding authors upon reasonable request.

## Acknowledgements

We thank K. Ogomori and C. Harada for technical assistance. This work was supported by grants from the Platform for Drug Discovery, Informatics and Structural Life Science by the Ministry of Education, Culture, Sports, Science and Technology (MEXT), and JSPS KAKENHI grants 21H05037 (O.N.), 22K19371 and 22H02751 (W.S.), and 21J20692 (T.T.); ONO Medical Research Foundation (W.S.); The Kao Foundation for Arts and Sciences (W.S.); The Takeda Science Foundation (W.S.); The Uehara Memorial Foundation (W.S.); the Platform Project for Supporting Drug Discovery and Life Science Research (Basis for Supporting Innovative Drug Discovery and Life Science Research (BINDS)) from AMED, under grant numbers JP19am01011115 (support no. 1109, O.N.).

## Author contributions

A. I. performed all the experiments. F.S., H.A., and H.S.O assisted with the cryo-EM data collection and the single particle analysis. W.S. designed the experiments. The manuscript was mainly prepared by A. I. and W.S., with assistance from O.N.

