## Supplementary Figure for "Structure and dynamics of the RF-amide QRFP receptor GPR103"

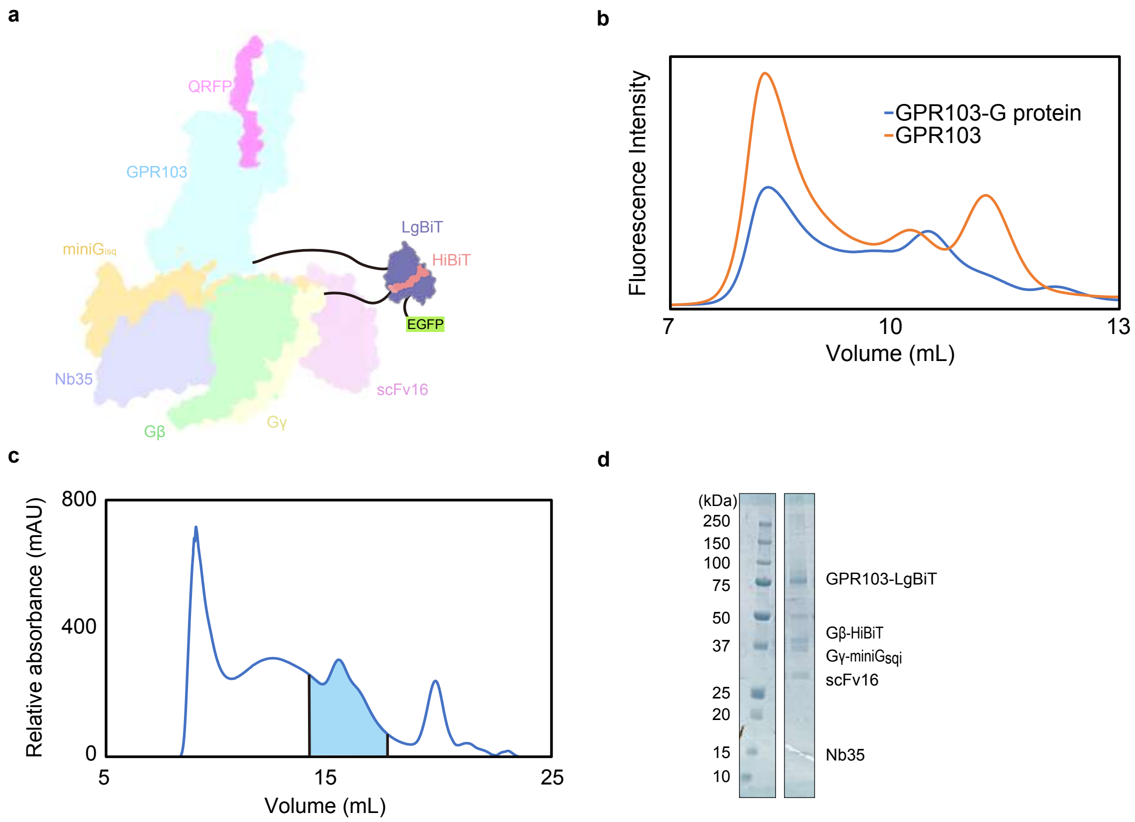


**Supplementary Figure 1 | Sumple purification.**
**a** Schematic representations of the fusion-G system. **b** Fluorescence-detection size-exclusion chromatography (FSEC) analysis of complex formation by the GPR103. Solubilised cells expressing only the GPR103 are orange, and co-expressing the GPR103 and G-protein are blue. **c** Size-exclusion chromatography of the GPR103-G-protein complex on Superose 6 increase column. The blue fraction was collected. **d** SDS-PAGE gel of samples after Size-exclusion chromatography.

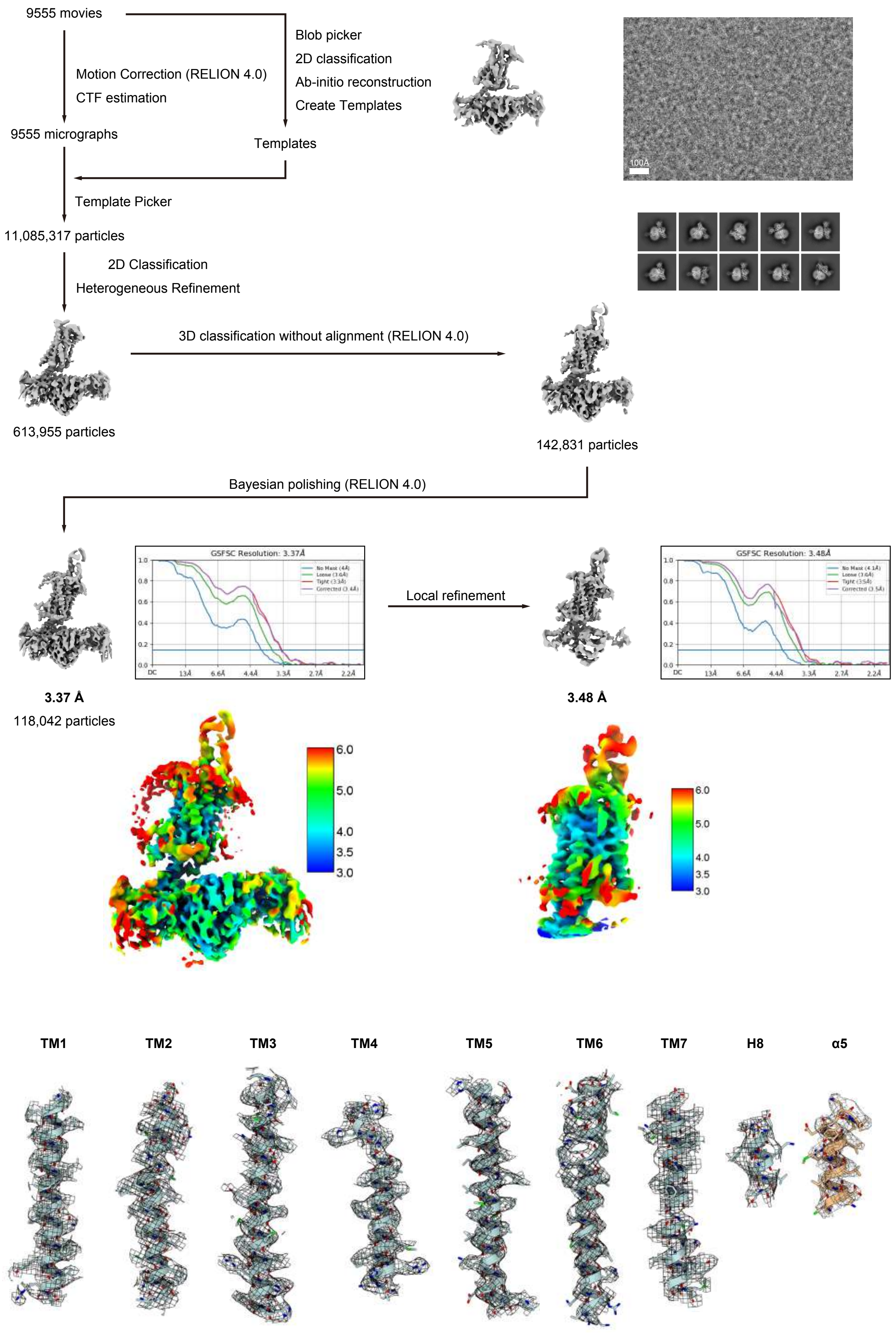


**Supplementary Figure 2 | Cryo-EM.**
Flow chart of the cryo-EM data processing for the GPR103-miniG$_q$ complex, including particle projection selection, classification, and 3D density map reconstruction. The 3D density map was refined with a mask on the receptor. Details are provided in the Methods section.

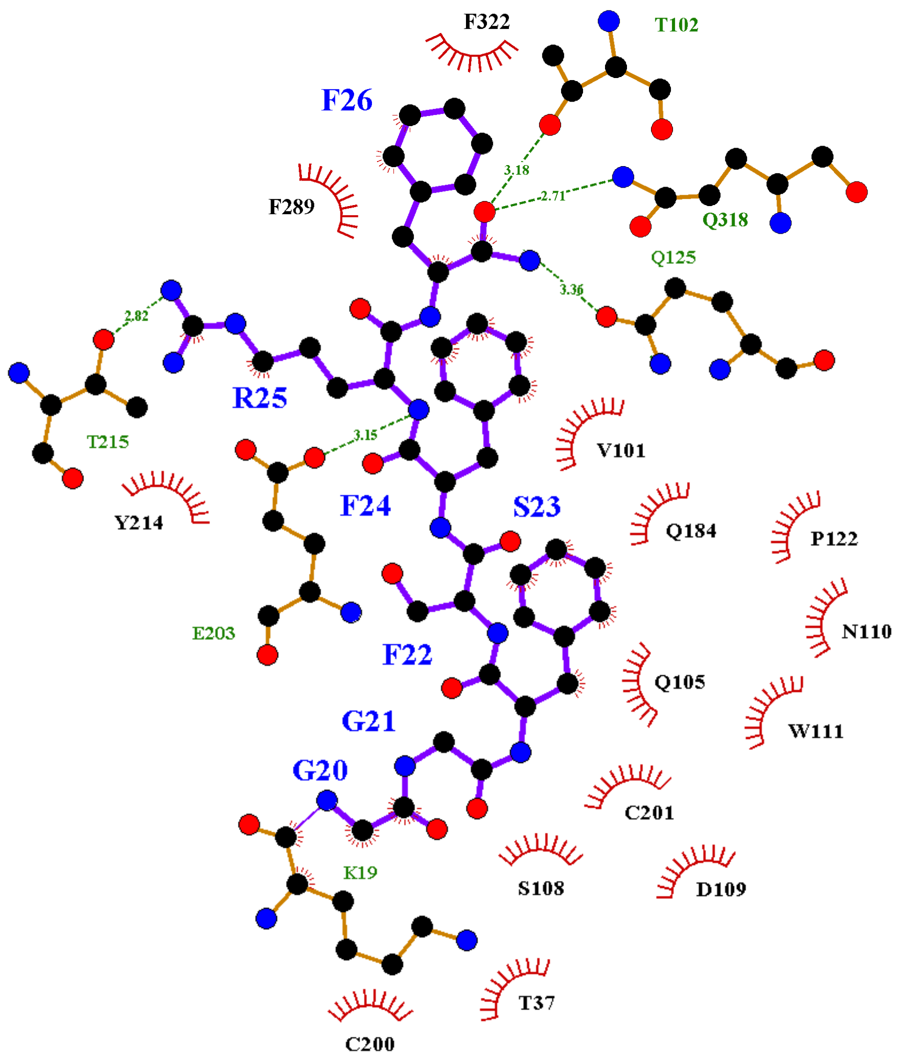


**Supplementary Figure 3 | Binding site diagram.**
Residues within 4.5 Å of the ligand are analysed by LIGPROT.

**a**

```
                     1         10          20         30         40         50
Human       .......MVRPYPLIYFLFLPLGACFPLLDRREPTDAMGGLGAGERWADLAMGPRPHSVW
Rat         ........MRCLCSWLCLLLPLSACFPLLDRRGPTD.IGDIGARMSWVQLTEGHTPRSV.
Chicken     .......MRAPYSLTCLILLSLGACCPPHEHHEPRHPEDVL..E...........PCWRG
Zebrafinch  .......MRAPYSLSCLFLLSLGACIPPGERWEPAEPGEGAALGGGWQRAAEGRGPGWRA
Xenopus     .......MRGSYGMSVLVLLSLGYTFALQDSREQSDPWERIRLLRMMADG.EENSAGALW
Zebrafish   MKIQVFHLSSTLQTTVFFLLVL.LVHPPRSLMLPHQPMVYLP.........RLYNPEWEA

                                                 60          70
Human       ..............................GS..SRWLRASQPQ.ALLVI....ARG.LQ
Rat         ......................................QSPRPQ.ALLVV....AKE.QQ
Chicken     AVA...........EAVGPCG.........WAA.PMRRRSEELG.ALLGI....ARV.LR
Zebrafinch  GARRRRSEELDELLSITRGPG.........WRAGARRRRSEELE.ALLSI....ARE.LR
Xenopus     ..............................YPLAPRQRKSTDPA.SLFSV....AKE.LQ
Zebrafish   AMLRL........QASLGAPGGRPVADLQPWALFQD.LGSEELLDRVNEILERQQRDIVK

                      QRFP43 ____________________________________________
                                          QRFP26 _________________________
                 80         90        100        110            120
Human       TSGREHAGCRFRFGRQDEGSEATGFLPAAG..EKTSG.....PLGNLAEELNGYSRKKGG
Rat         ASRREHTG..FRLGRQDSGSEATGFLPTDS..EKASG.....PLGTLAEELSSYSRRKGG
Chicken     GYGQQHSVG.............PRGRPEGSEKRGGGG.....TLGDLAEELNGYGRKKGG
Zebrafinch  GYSAAGAGQR......PGGSGGPGALPVVG..EKRSG.....TLGNLAEEINGYNRRKGG
Xenopus     GFGKERAGFRFRFGRQEEGNEFEDFEQQDE..EKRGG....TALGSLAEELNGYNRKKGG
Zebrafish   GQKRE.ELERRPMGTFPDNHIMPLLYPQGGEAEEVGGEKRNEALTSIAGGLQAFNRQKGG

            _____
            _____
             130
Human       FSFRF
Rat         FSFRF
Chicken     FAFRF
Zebrafinch  FTFRF
Xenopus     FSFRF
Zebrafish   FGFRF
```

**b**

| No. | 37 | 98 | 101 | 102 | 104 | 105 | 108 | 109 | 110 | 111 | 118 | 122 | 125 | 126 | 129 |
|---|---|---|---|---|---|---|---|---|---|---|---|---|---|---|---|
| BW | Nterm | 2.57 | 2.6 | 2.61 | 2.63 | 2.64 | 2.67 | 2.68 | 23.49 | 23.50 | 3.25 | 3.29 | 3.32 | 3.33 | 3.36 |
| Human | T | C | V | T | L | Q | S | D | N | W | C | P | Q | S | V |
| Rat | T | C | V | T | L | Q | S | D | K | W | C | P | Q | S | V |
| Chicken | V | C | F | T | L | Q | S | S | E | W | C | P | Q | S | I |
| Zebrafinch | I | C | F | T | L | Q | S | S | N | W | C | P | Q | S | I |
| Xenopus | I | C | F | T | L | Q | S | S | N | W | C | P | Q | S | I |
| Zebrafish | I | C | F | T | L | Q | S | S | E | W | C | P | Q | T | V |

| No. | 180 | 184 | 199 | 200 | 201 | 202 | 203 | 205 | 211 | 214 | 215 | 218 | 222 | 286 | 289 | 293 | 297 | 318 | 322 |
|---|---|---|---|---|---|---|---|---|---|---|---|---|---|---|---|---|---|---|---|
| BW | 4.6 | 4.64 | ECL2 | ECL2 | ECL2 | ECL2 | ECL2 | ECL2 | 5.35 | 5.38 | 5.39 | 5.42 | 5.46 | 6.48 | 6.51 | 6.55 | 6.59 | 7.39 | 7.43 |
| Human | M | Q | I | C | C | L | E | W | Q | Y | T | I | L | W | F | H | E | Q | F |
| Rat | M | Q | I | C | C | L | E | W | Q | Y | S | I | L | W | F | H | E | Q | F |
| Chicken | M | Q | I | C | C | L | E | W | Q | Y | T | I | L | W | F | H | E | Q | F |
| Zebrafinch | M | Q | V | C | C | L | E | W | Q | Y | T | I | L | W | F | H | E | Q | F |
| Xenopus | M | Q | V | C | C | L | E | W | Q | Y | T | I | L | W | F | H | E | Q | F |
| Zebrafish | M | Q | V | C | C | Q | E | W | R | Y | A | I | L | W | F | H | E | Q | F |

**c**

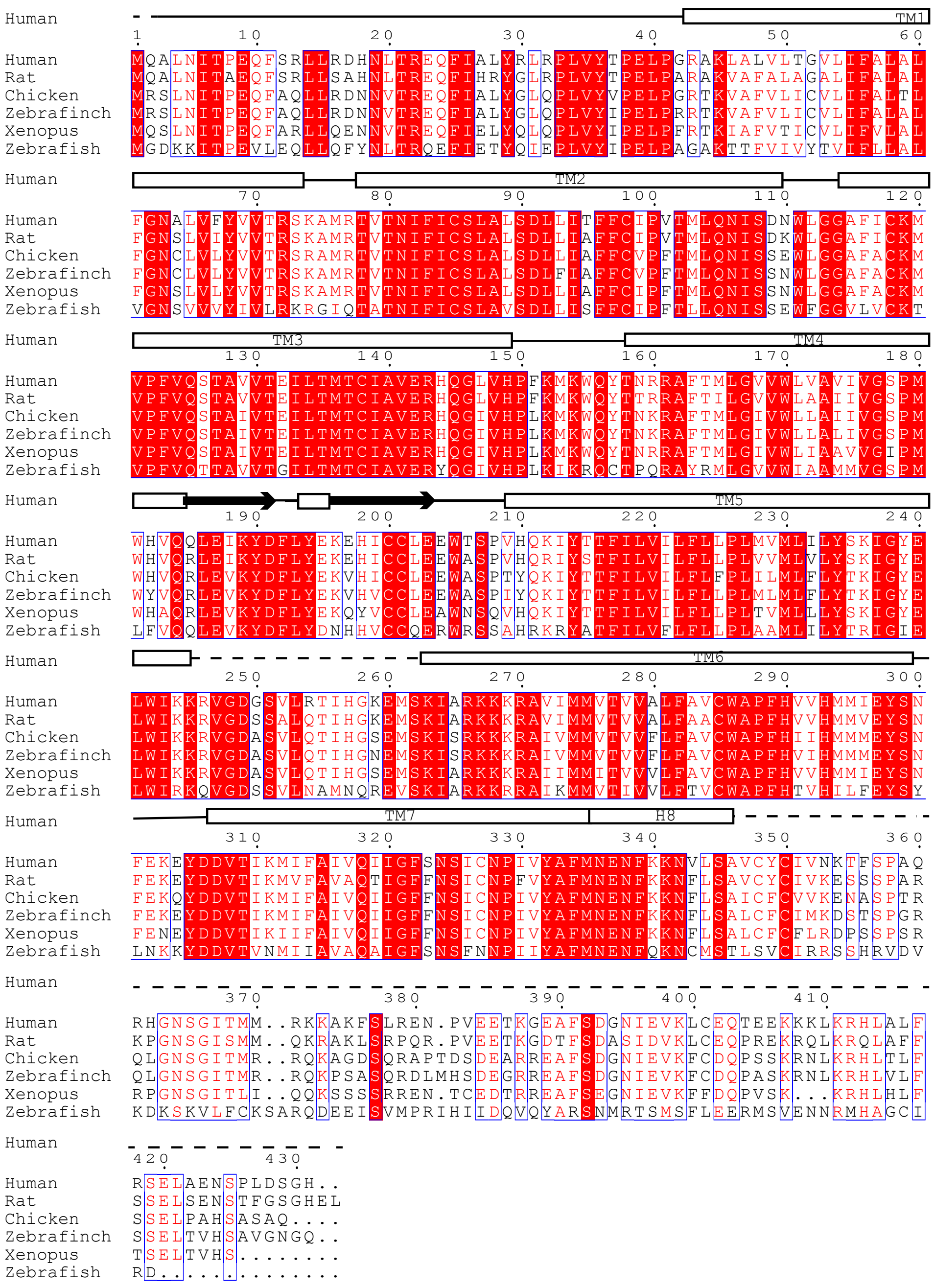


**Supplementary Figure 4 | Conservation of the interaction site.**
**a** Alignment of the amino acid sequences of QRFP homologs. **b** Comparison of residues within the TMD involved in QRFP binding in GPR103 homologues. **c A**lignment of the amino acid sequences of GPR103 homologs.

**a**

| | |
|---|---|
| QRFP | EDEGSEATGFLPAAGEKTSGPLGNLAEELNGYSRKKGGFSFRF |
| RFRP-1 | MPHSFANLPLRF |
| Kisspeptin-10 | YNWNSFGLRF |
| NPFF | FLFQPQRF |
| PrRP-31 | SRTHRHSMEIRTPDINPAWYASRGIRPVGRF |
| NPY | YPSKPDNPGEDAPAEDMARYYSALRHYINLITRQRY |
| CCK | KAPSGRMSIVKNLQNLDPSHRISDRDYMGWMDF |
| orexinB | RSGPPGLQGRLQRLLQASGNHAAGILTM |

**b**

| | | R25 | | | | | | | | |
|---|---|---|---|---|---|---|---|---|---|---|
| No. | | 180 | 203 | 205 | 211 | 214 | 215 | 218 | 293 | 297 |
| BW | | 4.6 | 45.52 | ECL2 | 5.35 | 5.38 | 5.39 | 5.42 | 6.55 | 6.59 |
| RF-amide receptors | GPR103 | M | E | W | Q | Y | T | I | H | E |
| | GPR10 | A | E | W | R | Y | A | L | N | D |
| | GPR54 | V | E | F | E | F | A | N | L | A |
| | GPR74 | S | E | W | R | Y | T | L | M | D |
| | GPR147 | S | E | W | R | Y | T | L | L | D |
| YRs | $Y_1$R | F | D | F | R | Y | T | L | N | D |
| | $Y_2$R | L | E | W | G | Y | S | S | Q | D |
| | $Y_3$R | F | E | W | R | Y | T | L | N | D |
| | $Y_4$R | F | E | W | R | Y | T | L | N | D |
| | $Y_5$R | L | E | W | R | F | T | L | H | D |
| OXRs | $OX_1$R | Q | E | N/A | P | Y | H | F | N | R |
| | $OX_2$R | Q | E | N/A | P | Y | H | F | N | R |
| CCKRs | $CCK_1$R | Y | E | N/A | Q | W | H | L | N | A |
| | $CCK_2$R | Y | H | N/A | R | W | S | L | N | A |

| | | F26 | | | | | | | | | | | | |
|---|---|---|---|---|---|---|---|---|---|---|---|---|---|---|
| | | | | | | | | | amide | | | | | |
| No. | | 126 | 129 | 214 | 218 | 222 | 286 | 289 | 125 | 322 | 98 | 105 | 102 | 318 |
| BW | | 3.33 | 3.36 | 5.38 | 5.42 | 5.46 | 6.48 | 6.51 | 3.32 | 7.43 | 2.57 | 2.64 | 2.61 | 7.39 |
| RF-amide receptors | GPR103 | S | V | Y | I | L | W | F | Q | F | C | Q | T | Q |
| | GPR10 | P | V | Y | L | T | W | L | Q | M | C | Y | T | H |
| | GPR54 | Q | V | F | N | L | W | I | Q | Y | C | L | T | H |
| | GPR74 | G | V | Y | L | I | W | L | Q | F | C | D | T | H |
| | GPR147 | G | V | Y | L | I | W | L | Q | F | C | D | T | H |
| YRs | $Y_1$R | C | I | Y | L | Q | W | L | Q | M | C | Y | T | H |
| | $Y_2$R | G | V | Y | S | L | W | L | Q | M | C | Y | T | H |
| | $Y_3$R | C | V | Y | L | Q | W | L | Q | M | C | D | T | H |
| | $Y_4$R | C | V | Y | L | Q | W | L | Q | M | C | Y | T | H |
| | $Y_5$R | C | V | F | L | Q | W | L | Q | M | C | S | T | H |
| OXRs | $OX_1$R | A | V | Y | F | T | Y | I | Q | Y | C | V | S | H |
| | $OX_2$R | T | V | Y | F | T | Y | I | Q | Y | C | V | T | H |
| CCKRs | $CCK_1$R | G | V | W | L | L | W | I | M | Y | C | P | N | L |
| | $CCK_2$R | G | V | W | L | L | W | V | M | Y | C | P | T | H |

**c**

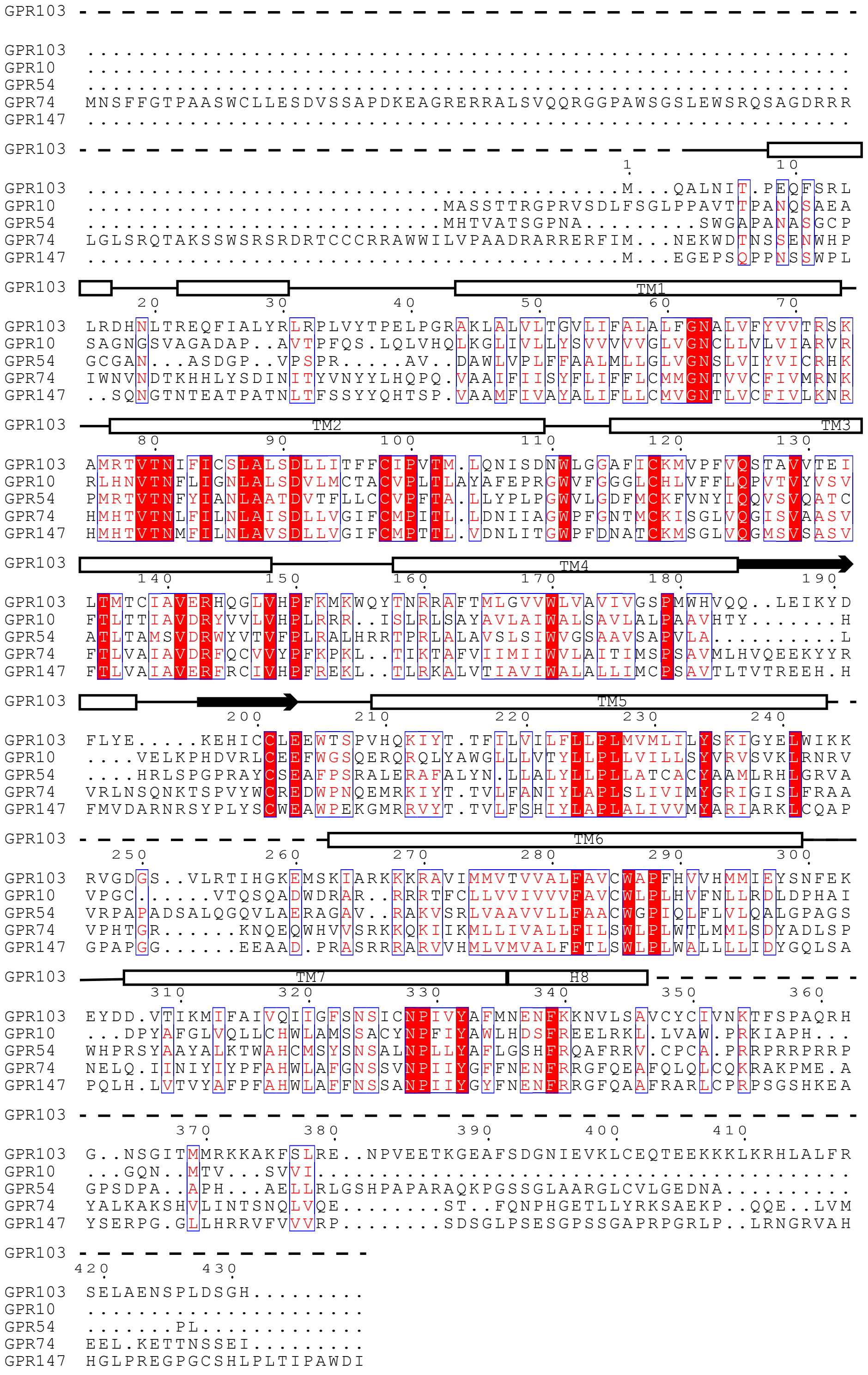


**Supplementary Figure 5 | Sequence comparison of GPR103-related peptide receptors.**
**a** Alignment of the amino acid sequences of GPR103-related peptides. **b** Comparison of residues interacting with GPR103 and RF-amide in RF-amide receptors, YR, CCKR and OXR. **c** Alignment of the amino acid sequences of RF amide receptors.

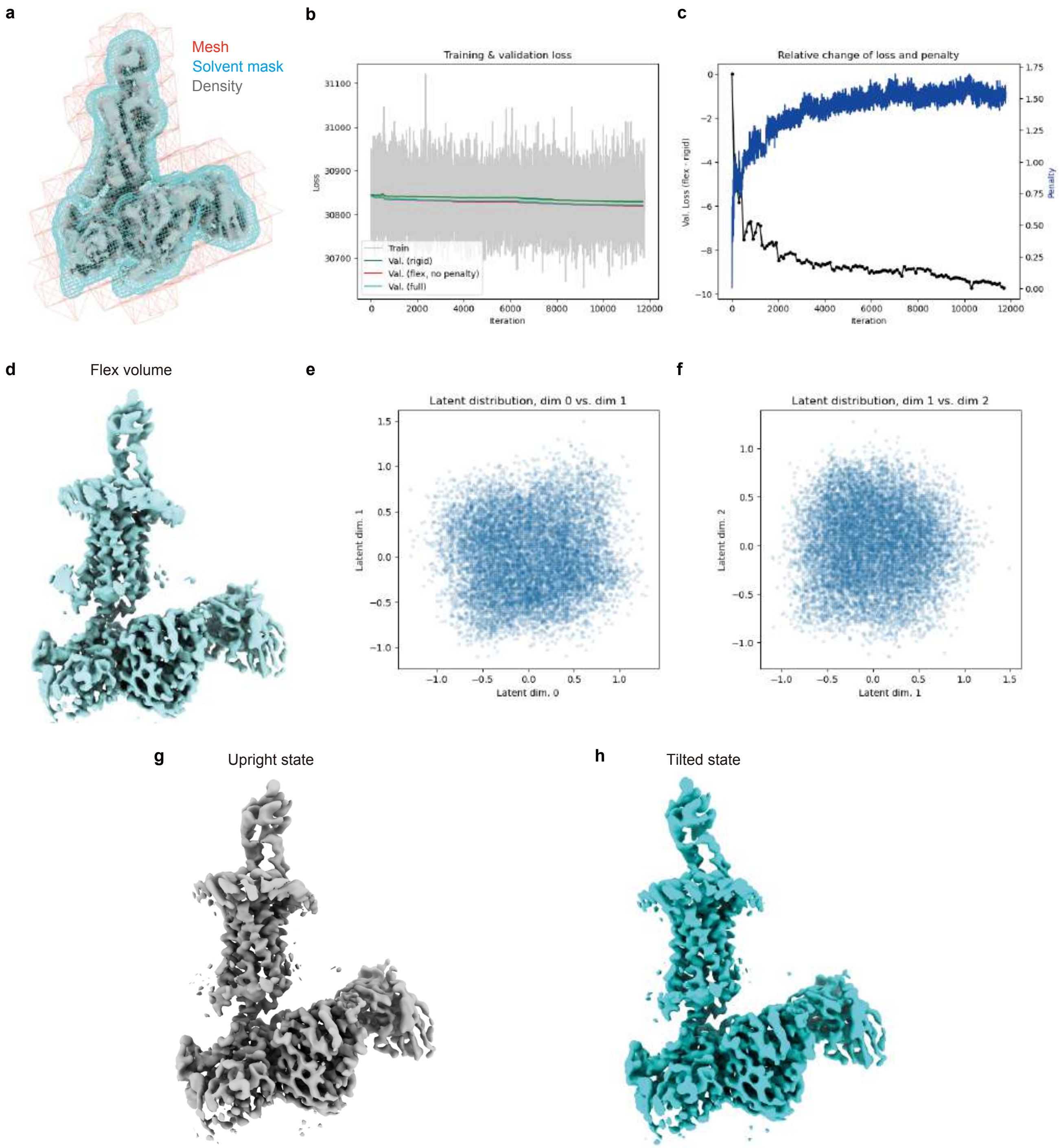


**Supplementary Figure 6 | 3DFlex refinement model.**
**a** A tetrahedral mesh, solvent mask and corresponding density which are used for 3DFlex. **b** Value of loss for each iteration. **c** In model validation, improvements in 3DFlex refinement compared to the non-3DFlex map through training iterations are shown in black, and changes in measured penalty compared to the unmodified map are shown in blue. **d** Flex volume: an output of high resolution reconstruction for 3DFlex. **e, f** Latent distribution of each dimensions describing the conformational heterogeneity in the particle set. **g, h** Representative of the volume series output of the 3DFlex generator job. Of the 40 classes corresponding to the largest eigenvalues, frame0 and frame40 were defined as the upright and tilted states, respectively.d cpk models.

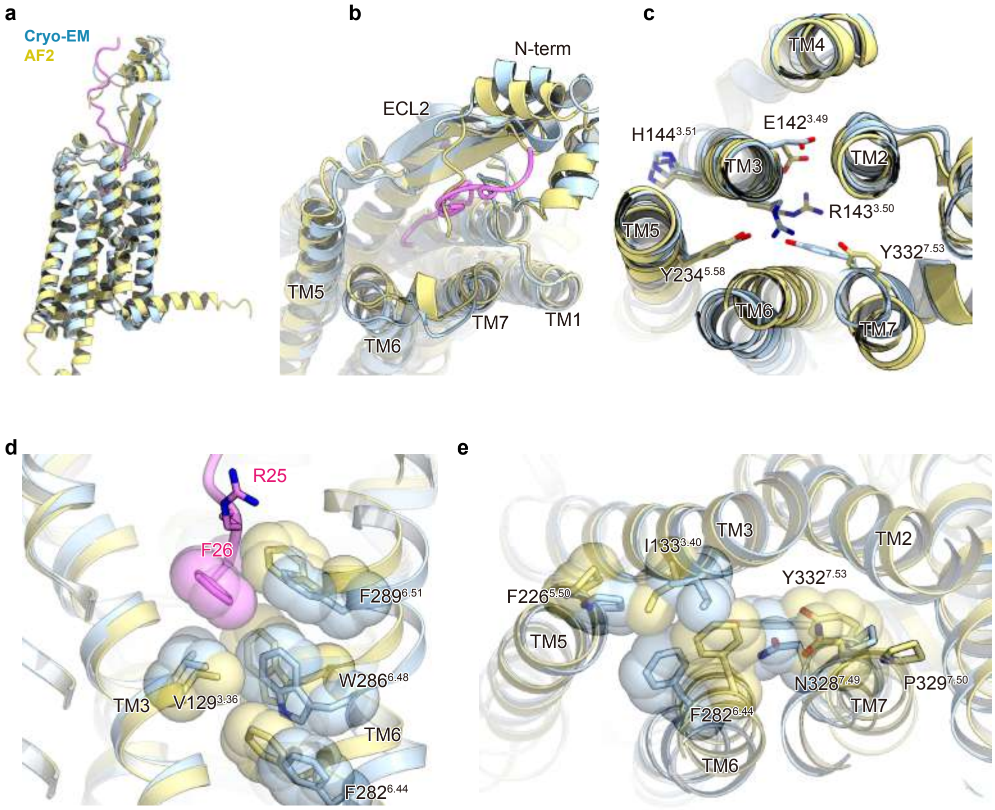


**Supplementary Figure 7 | Comparison with AF2 model.**
**a-e** Superimposition of the cryo-EM (blue) and AF2 (khaki) structures , **(a)** overall view of the receptor, **(b)** focused on the extracelluar side, and **(c)** focused on the intracelluar side. Critical activation motifs are shown by stick models. **(d)** Peptide-W$^{6.48}$ interactions, and **(e)** PIF and NPxxY motifs. The important residues are shown by stick and cpk models.
